# A reference assembly for the legume cover crop, hairy vetch (*Vicia villosa*)

**DOI:** 10.1101/2023.03.28.534423

**Authors:** Tyson Fuller, Lisa M. Koch, Lisa Kissing Kucek, Shahjahan Ali, Hayley Mangelson, Timothy Hernandez, Timothy P.L. Smith, Derek M. Bickhart, Heathcliffe Riday, Michael L. Sullivan

**Affiliations:** US Dairy Forage Research Center, United States Department of Agriculture Agricultural Research Service (USDA-ARS), 1925 Linden Drive, Madison, WI 53706 USA; Phase Genomics, 1617 8th Ave N, Seattle, WA 98109, USA; Noble Research Institute, 2510 Sam Noble Parkway, Ardmore, OK 73401, USA; US Meat Animal Research Center, United States Department of Agriculture Agricultural Research Service (USDA-ARS), PO Box 166 (State Spur 18D), Clay Center, NE 68933, USA

## Abstract

*Vicia villosa* is an incompletely domesticated annual legume of the Fabaceae family native to Europe and Western Asia. *V. villosa* is widely used as a cover crop and as a forage due to its ability to withstand harsh winters. A reference-quality genome assembly (Vvill1.0) was prepared from low error rate long sequence reads to improve genetic-based trait selection of this species. The Vvill1.0 assembly includes seven scaffolds corresponding to the seven estimated linkage groups and comprising approximately 68% of the total genome size of 2.03 gigabase pairs (Gbp). This assembly is expected to be a useful resource for genetic improvement of this emerging cover crop species as well as to provide useful insights into plant genome evolution.

## DATA DESCRIPTION

### Background

*Vicia villosa* Roth (hairy vetch) is a mostly outcrossing hermaphroditic diploid (2n = 2x = 14) annual legume originating from Europe and Western Asia [1,2]. *V. villosa* belongs to the *Vicia* genus of the Fabaceae family and is the second most cultivated vetch worldwide, used both as a forage species and as a cover crop [1,3,4]. *V. villosa* is especially useful as a winter cover crop for warm season crops (i.e. corn and soybeans) since it is one of few legumes that can survive in harsh winter conditions [5].

*V. villosa’s* use as a cover crop benefits cash crops primarily through nitrogen fixation, soil and water conservation, and its ability to produce biomass in a short period of time [3–5]. *V. villosa* is an incompletely domesticated species. Variations in pod dehiscence and seed dormancy across populations can result in reduced yields and increased weediness [6,7], which limits the adoption of *V. villosa* use by farmers [7,8].

Differences in chromosome number between species of the *Vicia* genus have been identified, making it an interesting model for studies of the plant genome [2,9,10]. Generation of reference genomes for species within the *Vicia* genus can be used to better understand the phylogeny and karyotype evolution of different species within the genus. Species-specific reference genomes can also aid in the identification of genes involved in beneficial and undesirable traits, ultimately increasing their use as cover crops by farmers. However, the first chromosome-level genome assembly within the *Vicia* genus (*V. sativa*, common vetch) has only recently been published [11].

High heterozygosity presents a unique challenge for generation of high-quality genome assemblies with current assembly methods. Heterozygous regions result in both false duplications of sequences and less contiguous assemblies [12–15]. This adversely impacts the final assembly size as well as other downstream analyses such as gene prediction and functional annotation [12,15]. We circumvent these difficulties by applying low-error rate long-read sequencing along with both manual and automated curation to generate a high-quality reference genome for the highly heterozygous *V. villosa*.

### Context

We present a high-quality reference genome assembly for *V. villosa*, representing only the second reference quality genome assembly in the *Vicia* genus. The assembly was compared with genome assemblies of other legume species including *V. sativa*. We observed a markedly higher level of heterozygosity in *V. villosa* compared to the selfing species, *V. sativa,* and demonstrate that the *V. sativa* reference is unsuitable as a proxy for variant calling with *V. villosa* DNA sequence data despite their common lineage. Our assembly, Vvill1.0, represents a reference-quality genomics resource for a common cover crop species, and provides further evolutionary insights into a unique clade of leguminous plant species.

## METHODS

### Sample information, nucleic acid extraction, and library preparation

A single individual was chosen from the ‘AU Merit’ [16] cultivar for its ability to be clonally propagated in tissue culture, and was named ‘HV-30.’ This individual of *V. villosa* was used for long-read and short-read DNA sequencing (Figure 1). Approximately 0.75 g of frozen leaf tissue from an individual plant was ground in a mortar and pestle under liquid nitrogen. High-molecular-weight DNA was extracted using the NucleoBond HMW DNA extraction kit as directed by the manufacturer (Macherey Nagel, Allentown, PA, USA). The DNA pellet was resuspended in 150 μL of 5 mM Tris-Cl pH 8.5 (kit buffer HE) by standing at 4 °C overnight, with integrity estimated by fluorescence measurement (Qubit, Thermo Fisher, Waltham, MA, USA), optical absorption spectra (DS-11, DeNovix, Willmington, DE, USA) and size profile (Fragment Analyzer, Thermo Fisher).

**Figure 1.**
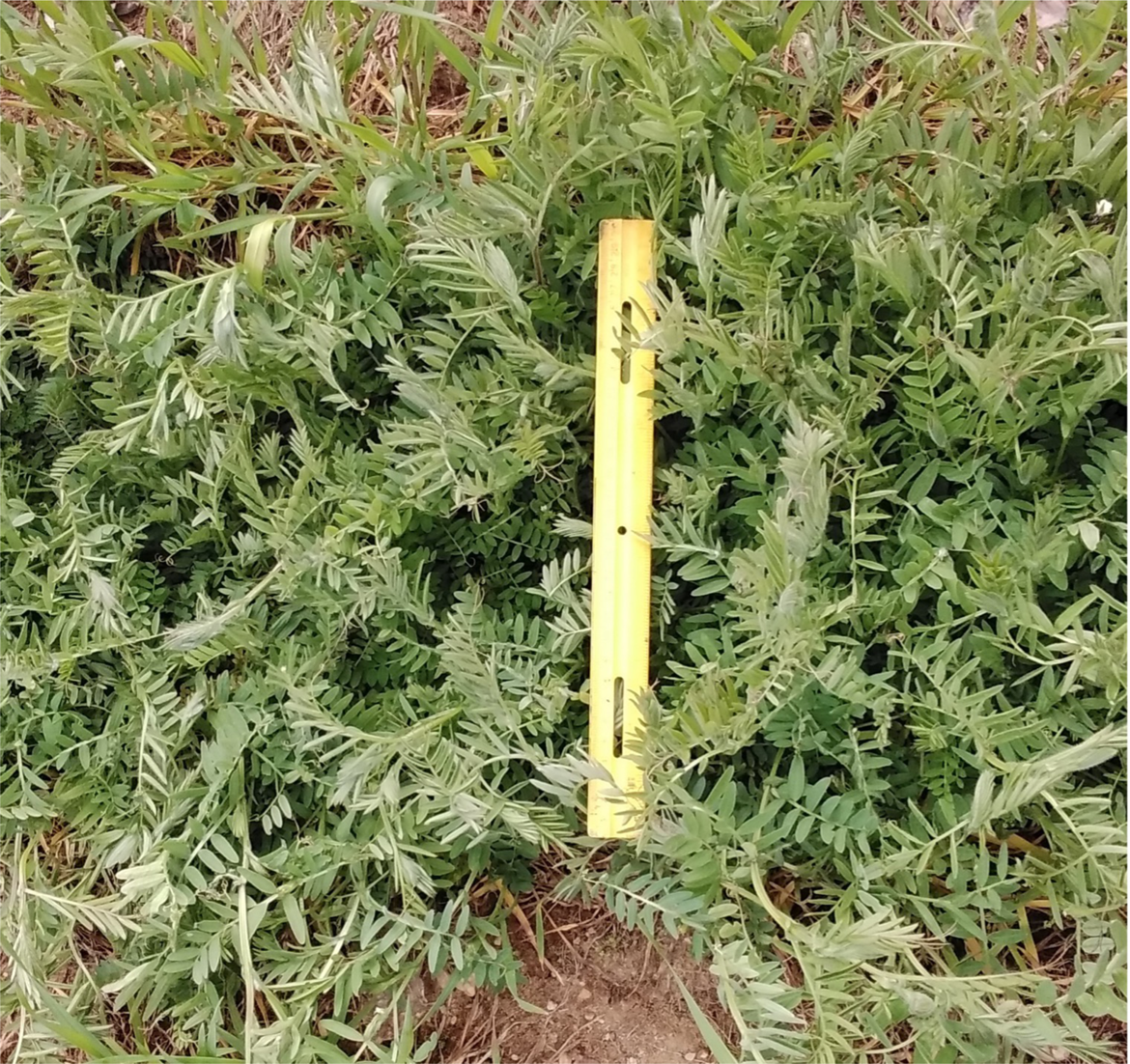
The HV-30 genotype of *Vicia villosa* was selected from the cultivar ‘AU Merit’ [16]. This image shows ‘AU Merit’ growing in Beltsville, MD on March 30^th^, 2022. A yellow 30 cm ruler is in the middle of the image for scale. The photo was taken by Allen Burke of USDA-ARS Beltsville Agricultural Research Center.

High molecular weight DNA used for high-fidelity long-read sequencing on the Pacific Biosciences (Menlo Park, CA, USA) Sequel II platform (HiFi sequence) was sheared (Hydroshear, Diagenode, Denville, NJ, USA) using a speed code setting of 13 to achieve a size distribution with peak at approximately 23 kbp. Smaller fragments were removed by size selection for >12 kbp fragments (BluePippin, Sage Science, Beverly, MA, USA). Size-selected DNA was used to prepare four SMRTbell libraries using the SMRTbell Express Template Prep Kit 2.0 as recommended by the manufacturer (Pacific Biosciences).

The DNA for short read sequencing was sheared to 550 bp on a Covaris M220 focused-ultrasonicator (Covaris, Woburn, MA, USA) by the University of Wisconsin-Madison Biotechnology Center (Madison, WI, USA) as specified in the TruSeq DNA PCR-Free Reference Guide (Illumina, San Diego, CA, USA) [17]. A library was prepared using 2 μg of the sheared DNA with the TruSeq DNA PCR-Free Library Preparation Kit, according to manufacturer guidance.

Total RNA was extracted from leaves, flowers, immature pods, seed coats, and cotyledons (seeds without seed coat) of HV-30 *V. villosa* plants using the Purelink RNA Mini Kit (Invitrogen, Carlsbad, CA, USA) with inclusion of the optional DNase treatment step. All tissue samples were frozen in liquid nitrogen immediately after collection and stored at -70 ^°^C until use. For leaves and flowers, 100 to 200 mg of tissue was ground in 2 mL prechilled microfuge tubes containing three 2.5 mm glass beads using a Mini-BeadBeater (BioSpec Inc., Bartlesville, OK, USA) at 5000 rpm for 1.75 min. RNA was extracted from the powdered tissue according to the manufacturer’s protocol with final elution of RNA in 100 μL RNase-free water.

Pod, seed coat, and cotyledon tissues (50-100, 100-200, and 25-50 mg, respectively) were ground with a mortar and pestle using liquid nitrogen. For pod and seed coat, the following modifications were made to the initial extraction: 2% (w/v) polyvinylpyrrolidone (PVP, average molecular weight 40,000) and 10% 2-mercaptoethanol (Sigma-Aldrich, St. Louis, MO) were added to the lysis buffer. Following centrifugation (12,000 x g, 5 min), one-half volume 3 M potassium acetate (pH 6.8) was added to the clear supernatant, the mixture incubated on ice for 30 min, and centrifuged at 12,000 x g for 30 min. The resulting supernatant was processed according to the PureLink protocol beginning with the addition of one-half volume of ethanol with final RNA elution in 100 and 30 μL RNase-free water for pod and seed coat, respectively. For cotyledon, 25 mg frozen powdered tissue was extracted using 1 mL Trizol reagent (Invitrogen, Carlsbad, CA, USA) in a 2 mL microcentrifuge tube. After phase extraction with 200 µl chloroform (Sigma-Aldrich) the clear upper phase was then processed according to the PureLink RNA protocol beginning with the addition of one-half volume of ethanol with final elution of RNA in 30 μL RNase-free water.

RNA concentrations were measured with broad or high sensitivity kits using Qubit 4.0 (Thermo Fisher) depending on RNA concentration of the samples. RNA quality was assessed by measuring RNA Integrity Number using a RNA IQ kit and Qubit 4.0. mRNA (PolyA+) was isolated from 100 ng of total RNA followed by RNAseq library preparation using TruSeq Stranded mRNA Library Preparation Kit (Illumina). The sequencing runs were performed on a HiSeq4000 (Illumina) with 2 x150 bp paired end (PE) read setting.

### Genome assembly and scaffolding

Genomic short-read libraries were sequenced on a NextSeq 500 instrument (Illumina) with a NextSeq High Output v2 300 Cycle Kit, generating 982 million 2 x150 PE reads. This resulted in 147.81 gigabase pairs (Gbp) of genomic sequence. These reads were used to estimate the total assembly length and heterozygosity of the sequenced *V. villosa* genotype. A histogram of abundance of 21-base length k-mers derived from the reads was generated from *V. villosa* short-read data using the Jellyfish version 2.2.9 tool [18] and was uploaded to the GenomeScope tool (http://qb.cshl.edu/genomescope/) [19] which estimated the haploid genome size to be 883 megabase pairs (Mbp). The expected genome size of 2.0 Gbp [20] is much larger, but k-mer-based estimations are generally underestimates. The estimated heterozygosity of *V. villosa* was 3.14% (Figure 2), which was substantially higher than that reported for *V. sativa* (0.09%) [11]. High degrees of heterozygosity present a substantial challenge for genome assembly with higher error-rate long-reads since errors and allelic variation are indistinguishable [21]. To circumvent this issue, low-error long-reads were used as the primary vehicle for genome assembly. A total of six SMRT cells were used with average insert length of 16.7 kilobase pairs (kbp), generating a total of 85.8 Gbp of sequence after processing for HiFi reads using the SMRT Link version 9.0 software with default settings (Pacific Biosciences). *V. villosa* primary contigs were generated using the PacBio IPA version 1.3.1 assembler and haplotigs were then screened for additional heterozygous duplications with purge_dups version 1.0.1 (https://github.com/dfguan/purge_dups) which identified 54 Mbp of duplicated sequence [22]. All duplicated sequence was removed from the primary haplotig assembly prior to scaffolding resulting in 5373 contigs with an N50 of approximately 600 kbp (Table 1).

**Figure 2.**
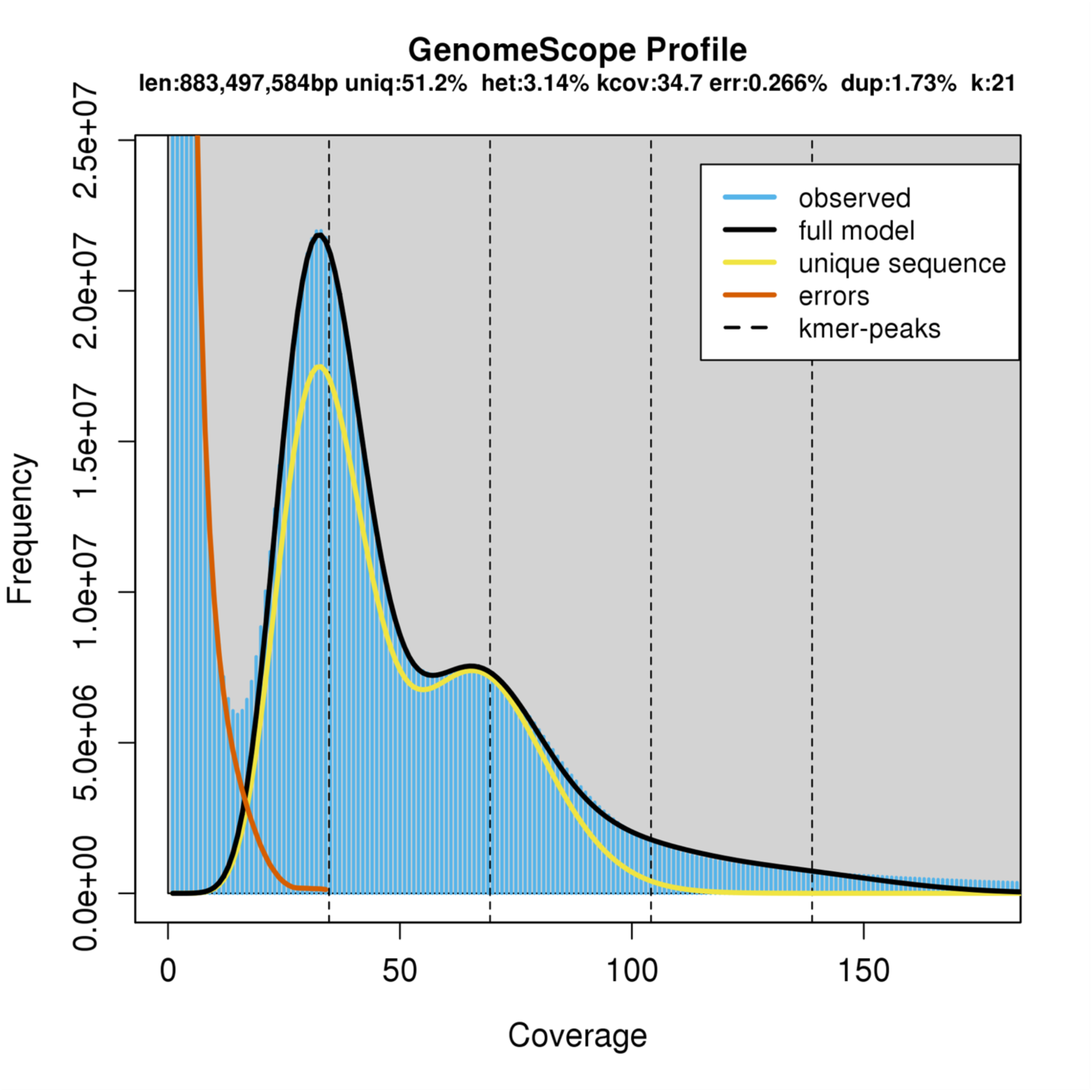
GenomeScope k-mer profile of *V. villosa* short-read data.

**Table 1.** Overview of *Vicia villosa* genome assembly.

| Feature | Value |
| --- | --- |
| Assembly size (bp) | 2,034,988,938 |
| No. of scaffolds | 1,888 |
| No. of contigs | 5,373 |
| Contig N50 | 604,665 |
| Scaffold N50 | 174,244,450 |
| GC content (%) | 35.62 |

Assembly scaffolding consisted of a combination of automated and manual processes. Chromatin conformation capture data was generated using a Phase Genomics (Seattle, WA) Proximo Hi-C 4.0 Kit, which is a commercially available version of the Hi-C protocol [23]. Intact cells from the sample were crosslinked using a formaldehyde solution as per the manufacturer’s supplied protocol, digested using a cocktail of restriction enzymes (DPNII, DdeI, HinfI, and MseI), end repaired with biotinylated nucleotides, and proximity ligated to create chimeric molecules composed of fragments from different regions of the genome that were physically proximal in vivo. Molecules were pulled down with streptavidin beads and processed into an Illumina-compatible sequencing library as recommended in the protocol. Sequencing was performed on an Illumina NovaSeq, generating a total of 140,472,036 2 x150 PE reads.

Reads were aligned to the primary haplotig assembly following the manufacturer’s recommendations [24]. Briefly, reads were aligned to the haplotig assembly using BWA-MEM [25] with the -5SP and -t 8 options specified, and all other options set to the default. SAMBLASTER [26] was used to flag PCR duplicates, which were later excluded from analysis. Alignments were then filtered with SAMtools [27] using the -F 2304 filtering flag to remove non-primary and secondary alignments. Putative misjoined contigs were broken using Juicebox [28,29] based on the Hi-C alignments. A total of 192 breaks were introduced, and the same alignment procedure was repeated from the beginning on the resulting corrected assembly.

Phase Genomics’ Proximo Hi-C genome scaffolding platform was used to create chromosome-scale scaffolds from the corrected assembly as described in Bickhart et al. [30]. As in the LACHESIS method [31], this process computes a contact frequency matrix from the aligned Hi-C read pairs, normalized by the number of restriction sites on each contig, and constructs scaffolds in such a way as to optimize expected contact frequency and other statistical patterns in Hi-C data. Approximately 60,000 separate Proximo runs were performed to optimize the number of scaffolds and scaffold construction in order to make the scaffolds as concordant with the observed Hi-C data as possible. Juicebox was used a second time to correct scaffolding errors.

Hi-C contact maps showed few off-diagonal contacts, which indicated agreement with the final scaffold structure (Figure 3). The few off-diagonal contacts present in the scaffold order are almost exclusively present on the telomeric ends of scaffolds, which indicates that they may be a biological signal from telomeric “bouquets” as opposed to scaffolding errors [32]. The final scaffolded assembly, Vvill1.0, represents the first reference quality genome assembly for a heterozygous out-crossing plant species in the *Vicia* genus to our knowledge [33].

**Figure 3.**
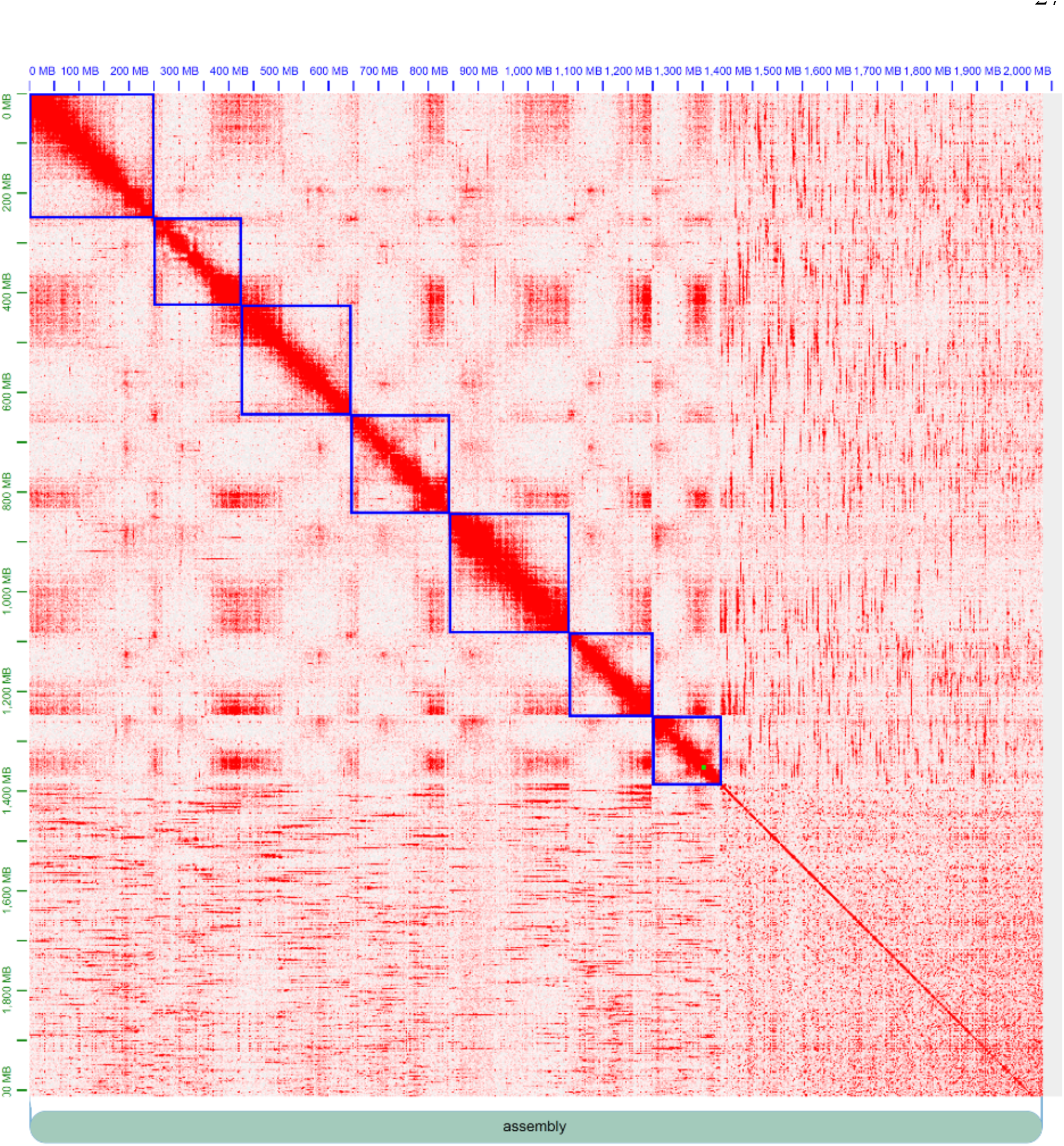
Hi-C link heatmaps and scaffold edits were produced by the JuiceBox tool [28]. Scaffold assignments (blue boxes) were identified from an optimal arrangement of signal along the diagonal.

The Vvill1.0 assembly was 2,034,988,938 bp total in 1,888 scaffolds, substantially larger than the GenomeScope haploid genome size estimate of 883 Mbp (Figure 2) but congruent with expectations from previous estimates [20]. The assembly had a scaffold N50 of 174.24 Mbp and a GC content of 35.62% (Table 1). Seven scaffolds of Vvill1.0 correspond to haploid representations of the seven estimated linkage groups for *V. villosa* [2] and comprise 67.74% of the total genome assembly size (Table 1) (Figure 4A). We note that a substantial proportion of the assembly (∼ 33% of base pairs; 1,881 scaffolds) was unable to be placed on distinct linkage group scaffolds due to the inherent heterozygosity of the individual. A combination of orthogonal genome assembly quality assessment tools was thus used to validate the completeness and accuracy of the assembly.

**Figure 4.**
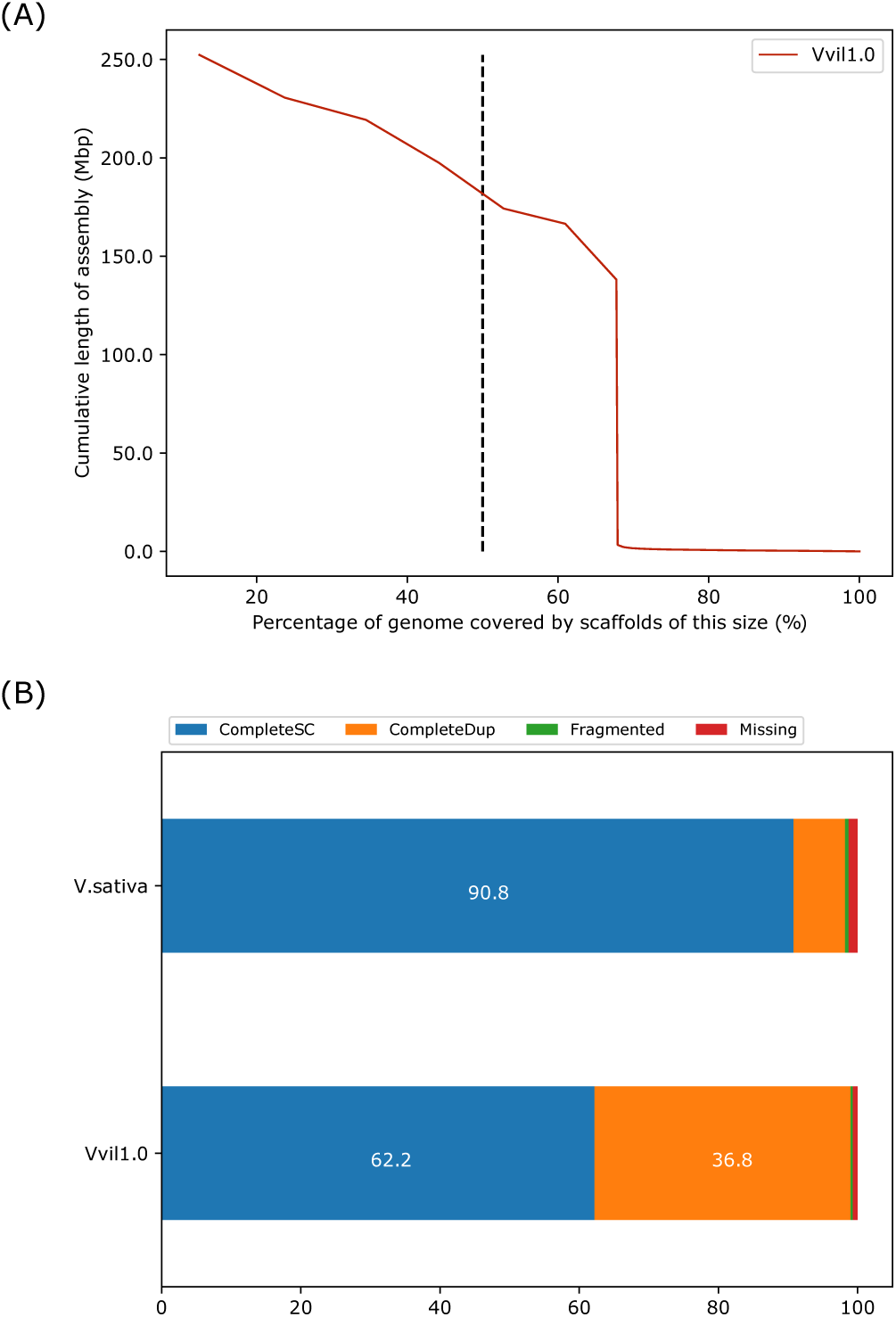
(A) Scaffold N(X) plot displaying the percentage of the genome (x-axis) covered by scaffolds of a specific length (y-axis). The vertical dotted line at the 50^th^ percentile of the genome length indicates the effective NG50 of the Vvill1.0 assembly. (B) The percentage of complete (CompleteSC), duplicated (CompleteDup), fragmented (Fragmented) and missing (Missing) single copy orthologous genes from Vvill1.0 and *V. sativa* (V.sativa) identified using the BUSCO [34] software package. The eudicots_odb10 data set (2326 markers) was used as the library for detecting single copy orthologs in both assemblies.

## DATA VALIDATION AND QUALITY CONTROL

All assembly validation and quality control data were produced by the Themis-ASM [35] pipeline run on the Vvill1.0 and *V. sativa* [11] genome assemblies with default settings. Briefly, the short-read dataset was aligned to each assembly using the BWA and SAMtools software packages [27,36]. Short-read alignments revealed that 98.6% of the *V. villosa* reads mapped to the Vvill1.0 assembly; however, only 47.0% of the *V. villosa* reads mapped to the *V. sativa* assembly. Similar comparisons using short-reads from *V. sativa* revealed a mapping rate of 64.0% and 99.7% to the Vvill1.0 and *V. sativa* reference assemblies, respectively, revealing a similar divergence in sequence profile in whole genome sequencing (WGS) read alignments. The *V. villosa* reads that did map to *V. sativa* had multiple single nucleotide variants and insertion-deletion mutations (INDELs), suggesting that frequent small variants may also cause issues with genome alignment comparisons despite the fact that the two species belong to the same genus. The frequency of sequence variants was confirmed by Freebayes analysis of short-read alignments [37]. Freebayes variant calls were used to generate a quality value (QV or Phred [38]) score for all bases with at least 3x coverage as described previously [30]. The base QV for our Vvill1.0 assembly was 45.02 indicating >99.99% accuracy of genome sequence when compared to short-read alignments (Table 2). Read alignments of *V. villosa* short-read data to the *V. sativa* reference produced a suboptimal QV of 14.66, which represents a difference in base alignment quality of three orders of magnitude compared to the Vvill1.0 assembly. We note that such comparative statistics are not indicative of any deficiency in the *V. sativa* assembly but reflect the advantages of a species-specific reference assembly for *V. villosa* genomic analyses.

**Table 2.**
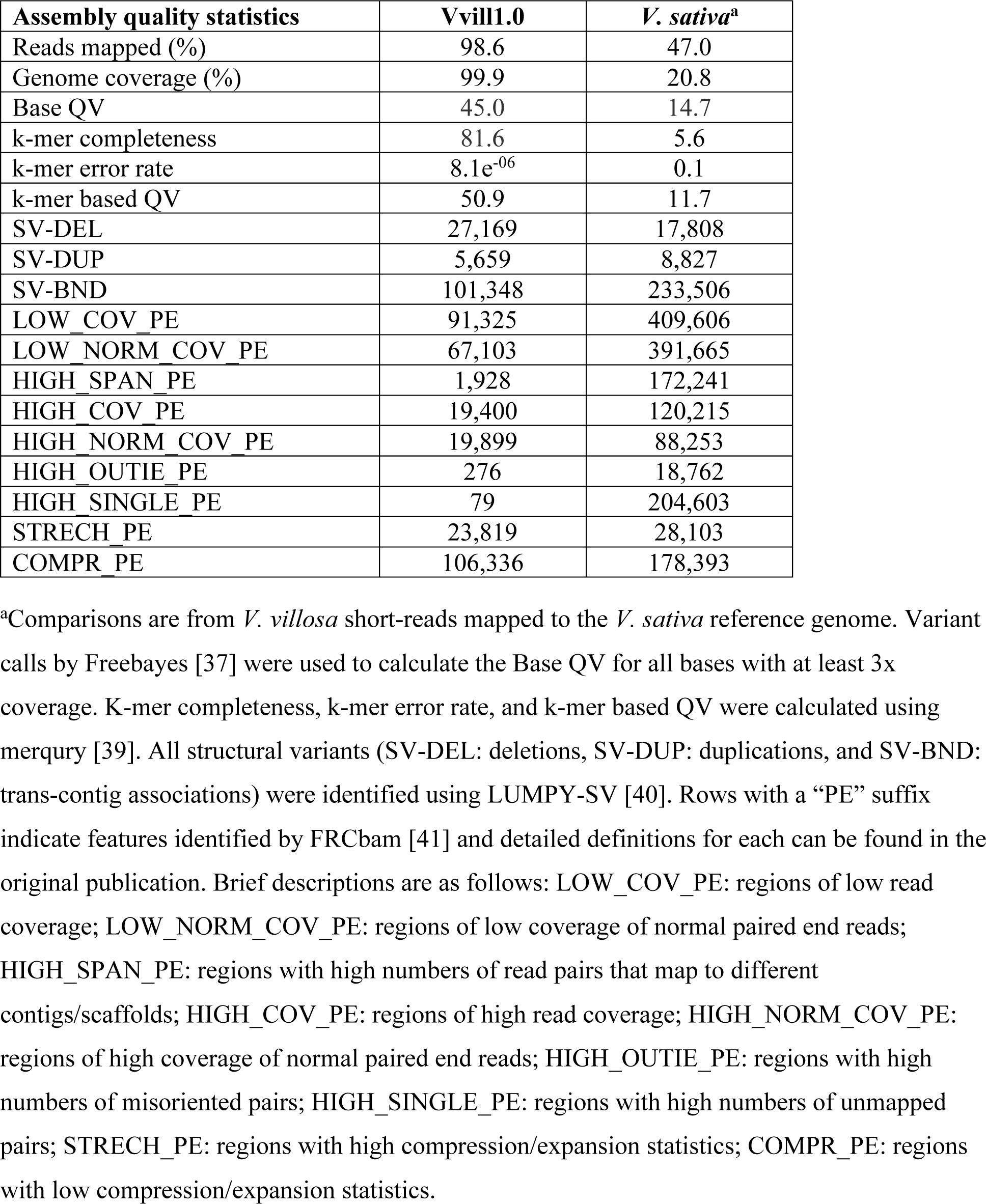
Read mapping statistics of Vvill1.0 and *V. sativa* genome assemblies using *V. villosa* short-reads.

| Assembly quality statistics | Vvill1.0 | *V. sativa*<sup>a</sup> |
| --- | --- | --- |
| Reads mapped (%) | 98.6 | 47.0 |
| Genome coverage (%) | 99.9 | 20.8 |
| Base QV | 45.0 | 14.7 |
| k-mer completeness | 81.6 | 5.6 |
| k-mer error rate | 8.1e<sup>-06</sup> | 0.1 |
| k-mer based QV | 50.9 | 11.7 |
| SV-DEL | 27,169 | 17,808 |
| SV-DUP | 5,659 | 8,827 |
| SV-BND | 101,348 | 233,506 |
| LOW_COV_PE | 91,325 | 409,606 |
| LOW_NORM_COV_PE | 67,103 | 391,665 |
| HIGH_SPAN_PE | 1,928 | 172,241 |
| HIGH_COV_PE | 19,400 | 120,215 |
| HIGH_NORM_COV_PE | 19,899 | 88,253 |
| HIGH_OUTIE_PE | 276 | 18,762 |
| HIGH_SINGLE_PE | 79 | 204,603 |
| STRECH_PE | 23,819 | 28,103 |
| COMPR_PE | 106,336 | 178,393 |

<sup>a</sup>Comparisons are from *V. villosa* short-reads mapped to the *V. sativa* reference genome. Variant calls by Freebayes [37] were used to calculate the Base QV for all bases with at least 3x coverage. K-mer completeness, k-mer error rate, and k-mer based QV were calculated using merqury [39]. All structural variants (SV-DEL: deletions, SV-DUP: duplications, and SV-BND: trans-contig associations) were identified using LUMPY-SV [40]. Rows with a “PE” suffix indicate features identified by FRCbam [41] and detailed definitions for each can be found in the original publication. Brief descriptions are as follows: LOW\_COV\_PE: regions of low read coverage; LOW\_NORM\_COV\_PE: regions of low coverage of normal paired end reads; HIGH\_SPAN\_PE: regions with high numbers of read pairs that map to different contigs/scaffolds; HIGH\_COV\_PE: regions of high read coverage; HIGH\_NORM\_COV\_PE: regions of high coverage of normal paired end reads; HIGH\_OUTIE\_PE: regions with high numbers of misoriented pairs; HIGH\_SINGLE\_PE: regions with high numbers of unmapped pairs; STRECH\_PE: regions with high compression/expansion statistics; COMPR\_PE: regions with low compression/expansion statistics.

The k-mer count plot for our assembly shows a large peak at ∼35x coverage representing k-mers from heterozygous sequences and a much smaller peak at ∼70x coverage representing k-mers from homozygous sequences (Figure 5). The approximately two-fold higher count of heterozygous compared to homozygous k-mers is in agreement with the high level of heterozygosity (3.1%) estimated by GenomeScope using the *V. villosa* short-reads as input (Figure 2). This elevated heterozygosity is likely a result of the cross-fertilizing nature of *V. Villosa*, compared with the selfing nature of *V. sativa* [33]. We note that the “read-only” k-mer peak, representing k-mers observed in the short-reads but not present in the assembly, indicates that some unique heterozygous sequence is not completely represented in Vvill1.0. This is likely a result of the removal of duplicated sequence resulting from the PacBio IPA assembly and the purge_dups workflow we used to generate Vvill1.0. The k-mer histogram plots are highly sensitive to the absence of single nucleotide variants that were likely present in purged duplicated regions, so their absence is less likely to impact future DNA sequence alignment surveys. This notable absence of k-mer frequency does provide a cautionary tale, as the purging of additional duplicated sequence would only exacerbate issues with genome representation as mentioned above.

**Figure 5.**
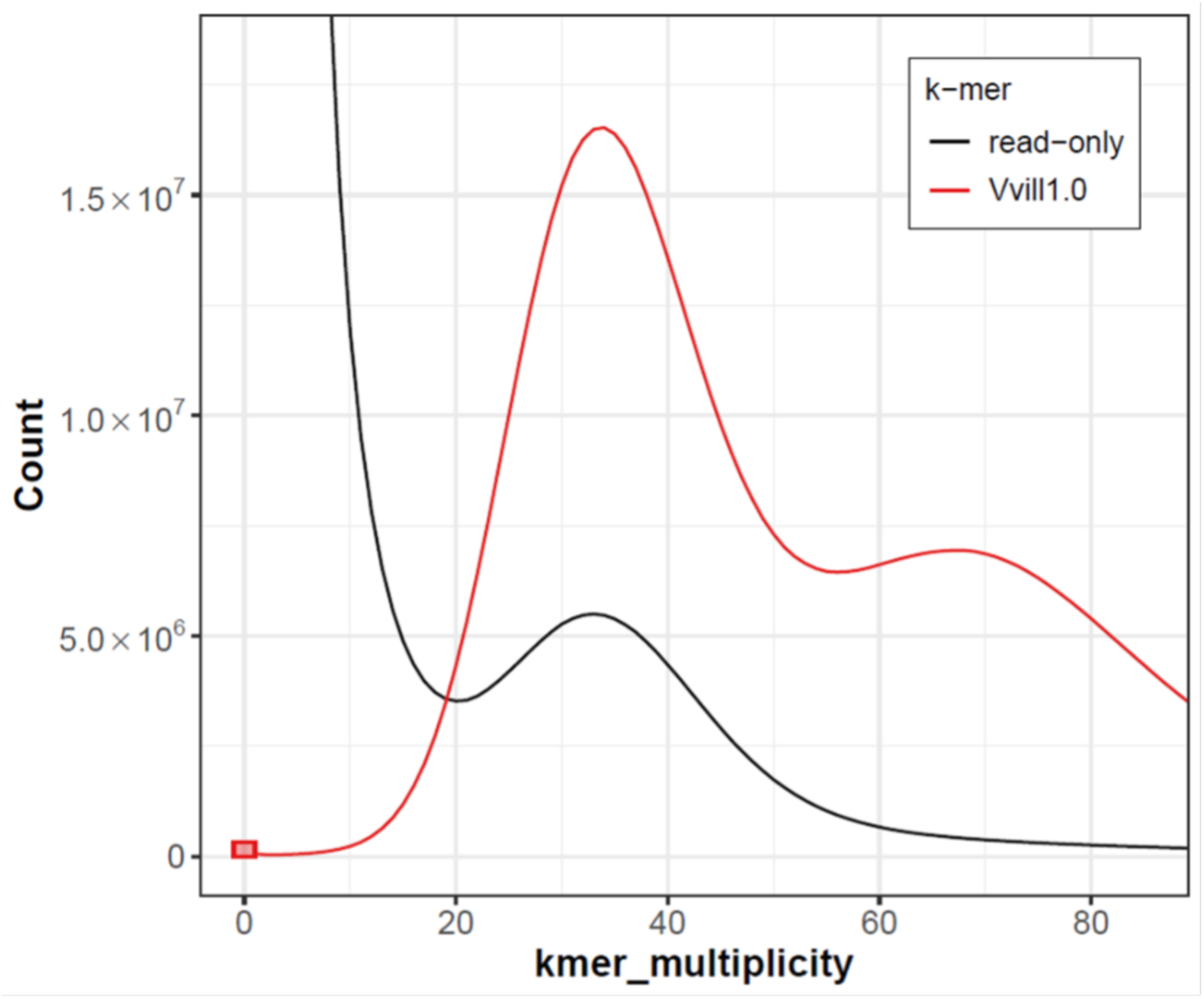
K-mer assembly spectra plot generated by merqury [39] showing the distribution of k-mers (k=21) found in the Illumina short-read set (black, read-only) and k-mers found in our Vvill1.0 assembly (red, Vvill1.0). The red bar at zero multiplicity indicates k-mers found only in the assembly. The read-only peak at ∼35x likely represents heterozygous variants missing from the assembly.

The discrepancies in alignment quality noted in our comparisons of *V. villosa* short-read data and the *V. sativa* reference assembly led us to question if there were significant structural discrepancies between the two species. The accuracy of structural variant prediction was assessed using LUMPY-SV [40] to call structural variants and FRCbam [41] to identify features or suspicious regions of the assembly based on read alignments, with *V. villosa* short-reads as input. The short-read alignments to the *V. sativa* genome assembly predicted 260,141 structural variants, with the majority predicted as complex structural variants (233,506). This is nearly twice as many structural variants predicted than when the same sequence reads were aligned to the *V. villosa* assembly (134,176). Further, the short-read alignments to the *V. sativa* genome had a substantially higher count of discordant genomic features than alignments to our *V. villosa* assembly (Table 2). These results suggest that smaller-scale (50-50,000 bp) structural variations in genome sequence exist between the two species.

Larger changes in genome structure were classified by identifying any candidate syntenic regions through whole-genome alignment. Minimap2 was used to identify pairwise alignments between our Vvill1.0 assembly and the *V. sativa* assembly using an alignment cutoff of 100,000 bp segments or greater (Figure 6) [42]. Some conserved segments of chromosomes were observed, but the majority of alignments are spread out between chromosome scaffolds of the two species. This variation in the genomic architecture suggests relaxed constraints on gene organization across these closely related species. By contrast, a similar whole genome alignment of two other legume species reference genomes shows better conservation of syntenic regions (Supplemental Figure 1). The chromosomal reorganization between these two species may underlie some of the phenotypic variation between them and further highlights the importance of having a species-specific genome reference assembly for future studies of wild vetch species.

**Figure 6.**
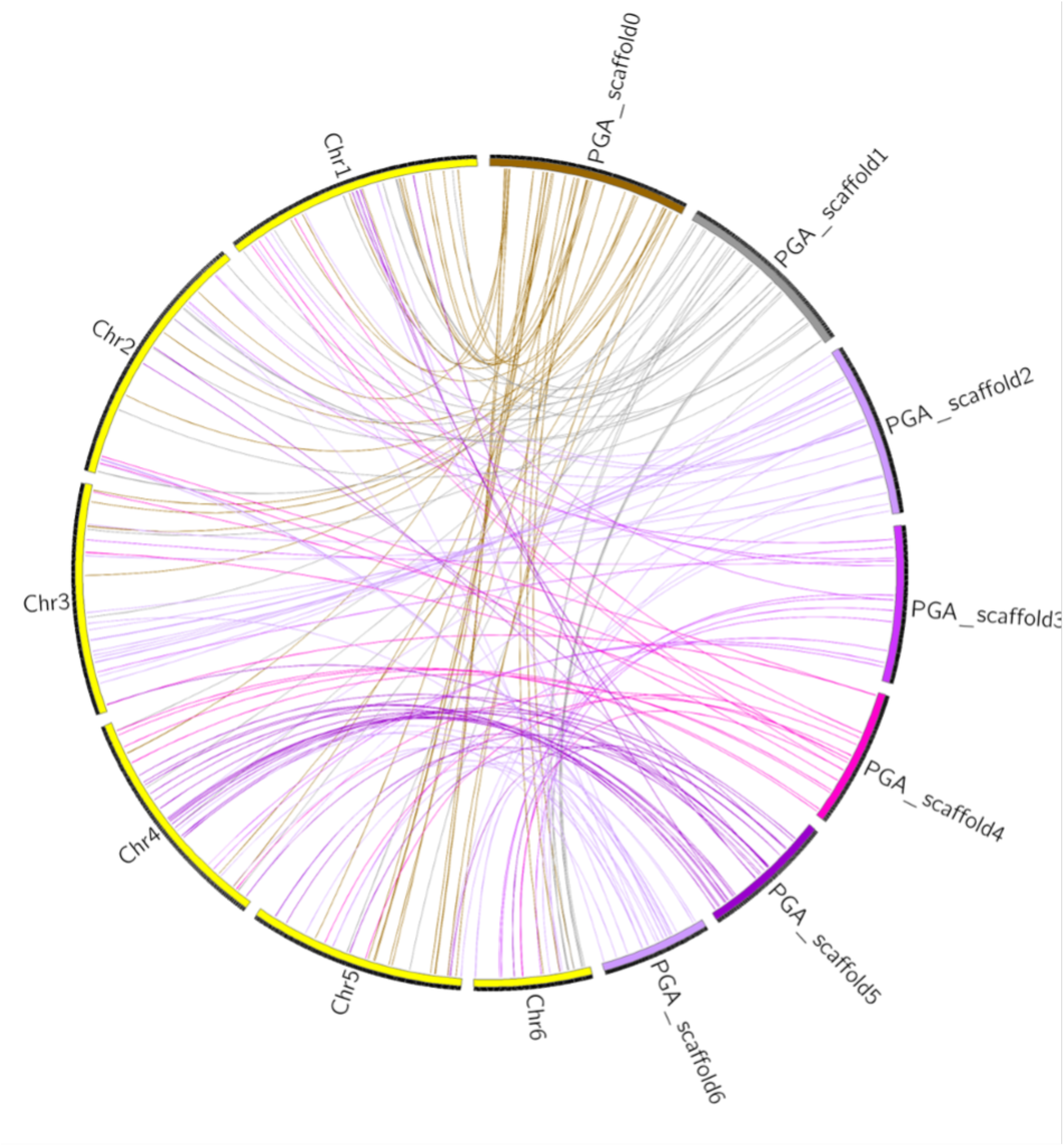
Circos plot showing syntenic regions shared between the *V. sativa* assembly (yellow outer bands) and Vvill1.0 (multi-colored outer bands) genomes [43]. Ribbons (colored matching the Vvill1.0 scaffolds) represent pairwise alignments of 100 kbp or larger identified using minimap2 [42].

### Genome annotation

Classification of all genic content and repetitive loci within Vvill1.0 was performed to increase its utility as a genomic resource. A list of canonical *V. villosa* repetitive elements was generated de novo using the EDTA version 2.0.0 software tool [44] with the “sensitive” setting to enable RepeatModeler (RRID:SCR_015027) recovery of transposable elements. The set of *V. villosa* canonical repetitive elements were then used as a custom library input to RepeatMasker version 4.0.6 [45], which was in turn used to soft-mask the Vvill1.0 assembly. Repetitive content was similar to the *V. sativa* reference assembly, with 81.1% of the assembly consisting of identified repeats in Vvill1.0 (Table 3), compared to 83.9% repetitive content in *V. sativa*. Comparisons of repetitive element lengths revealed few discrepancies in repeat content between the two vetch assemblies which had similar distributions of repeat fragment sizes for nearly all classes. A notable discrepancy was identified in the size distributions of miniature inverted-repeat transposable elements (MITE), where larger MITE_DTH and MITE_DTC elements were more prevalent in *V. villosa* and larger MITE_DTT elements were more prevalent in *V. sativa* (Figure 7). This suggests that differential expansion and amplification bursts of MITEs may have occurred in both lineages after their divergence.

**Table 3.** Repetitive element content of *V. villosa*.

| Repetitive elements |  | Number | Cumulative Length (bp) | Percentage of Genome |
| --- | --- | --- | --- | --- |
| Retroelements<sup>a</sup> |  | 1,080,921 | 830,932,491 | 60.0 |
|  | LINEs | 2,982 | 1,105,274 | 0.1 |
|  | LTRs | 1,077,939 | 829,827,217 | 59.9 |
| DNA Transposons |  | 802,725 | 224,578,692 | 16.2 |
| Unclassified |  | 221628 | 53995075 | 3.9 |
| Simple Repeats |  | 193714 | 11729117 | 0.9 |
| Low Complexity |  | 30795 | 1617938 | 0.1 |
| **Total** |  | **2,329,783** | **1,122,853,313** | **81.1** |

<sup>a</sup>Long interspersed nuclear elements (LINE), long terminal repeats (LTR)

**Figure 7.**
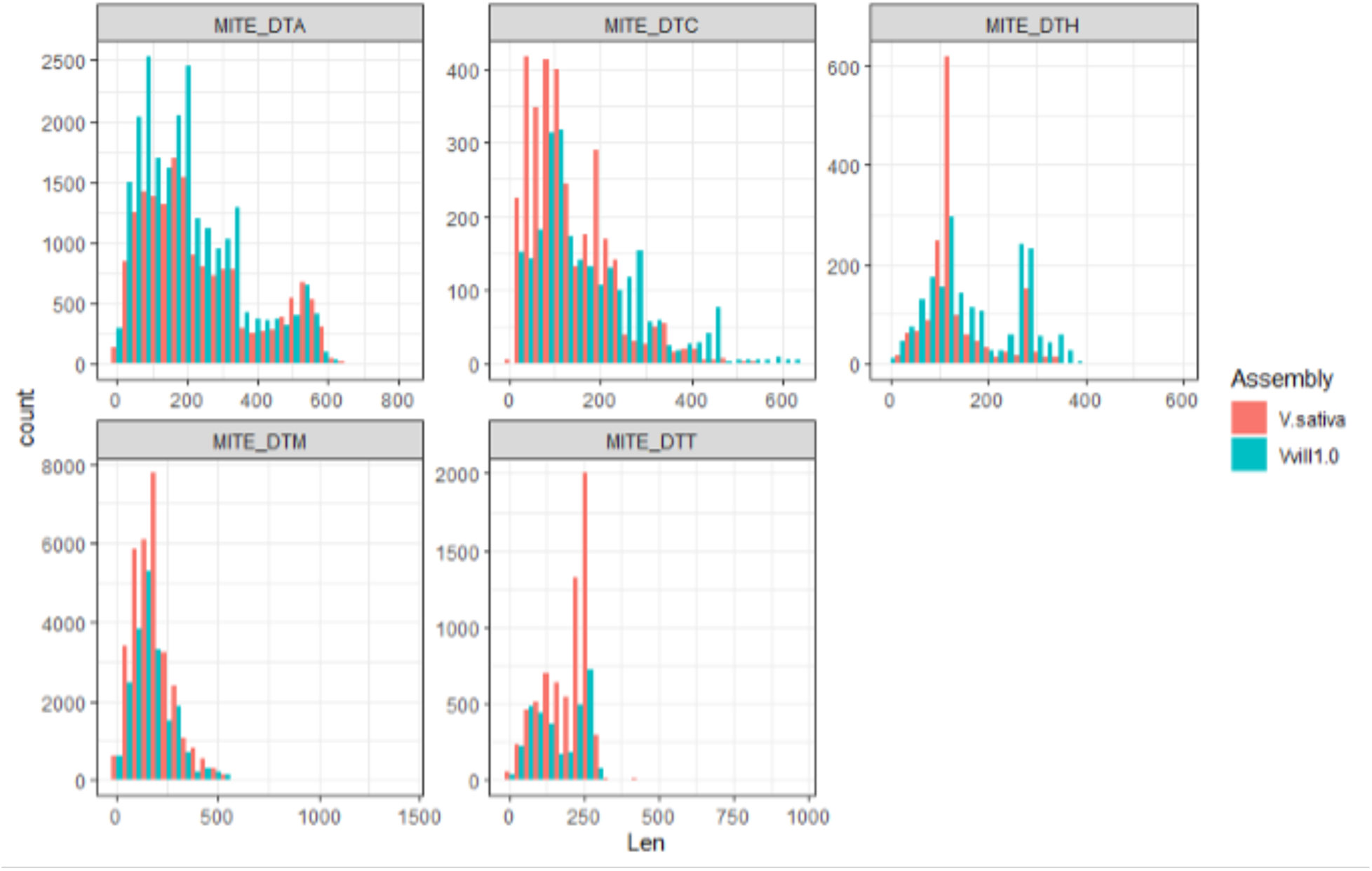
Length distribution of miniature inverted repeat transposable elements (MITE) repeats in *V. sativa* and *V. villosa.* MITE families are indicated by a suffix after the underscore in each subplot’s title, and follow Repbase (https://www.girinst.org/repbase/) naming classifications.

We annotated all coding sequences in the Vvill1.0 assembly using a combination of *ab initio* prediction and RNAseq evidence. RNAseq reads [citation to our GigaDB accession] were aligned to the soft-masked Vvill1.0 assembly using the STAR version 2.7.9 (RRID:SCR_004463) alignment tool in the “genomeGenerate” runtime mode. We identified 53,321 protein-coding genes (Table 4), which was nearly equivalent to the number of protein-coding genes (53,218) annotated in the *V. sativa* reference assembly.

**Table 4.** Gene annotation summary statistics.

| Features | Vvill1.0 | *V. sativa*<sup>a</sup> |
| --- | --- | --- |
| Protein-coding genes | 53,321 | 53,218 |
| Average exons per gene | 4.6 | 4.4 |
| Average exon length (bp) | 207.4 | 223.4 |
| Average intron length (bp) | 434.0 | 415.1 |

<sup>a</sup>Summary statistics for the *V. sativa* assembly were taken from [11].

Putative functions of identified coding sequences were identified through alignment of predicted protein amino acid sequence of *V. villosa* genes against the UniProt database release 2022_02 and the National Center for Biotechnology Information (NCBI) non-redundant database using the DIAMOND alignment tool version 2.0.14.152 [46]. The top scoring hit was chosen for each sequence (see GigaDB supplementary data files uniport_anno.tsv and ncbi-nr_anno.tsv for the DIAMOND output data for the UniProt and NCBI non-redundant databases, respectively.

Protein sequences were also aligned against the EggNOG database version 5.0.2 using EggNOGmapper version 2.1.8 in order to assign Kyoto Encyclopedia of Genes and Genomes (KEGG) pathways and KEGG orthologous groups (KO) to each sequence [47] (see GigaDB supplementary data file eggnog.tsv for the output data from EggNOGmapper). This allowed for the annotation of 43626 (81.8%) predicted protein-coding genes with at least one function (Table 5).

**Table 5.** Number of genes with functional annotations identified using different databases.

| Database |  | Number annotated | Percent annotated |
| --- | --- | --- | --- |
| NCBI-NR |  | 43455 | 81.5 |
| UniProt |  | 32445 | 60.9 |
| EggNOG | Pfam | 37949 | 71.2 |
|  | KEGG pathway | 12887 | 24.2 |
|  | KEGG KO | 20055 | 37.6 |
|  | GO | 20786 | 39.0 |
| **Total annotated** |  | 43626 | 81.8 |
| **Total** |  | 53312 |  |

### Phylogenetic tree construction

Large structural variations identified from *V. sativa* and *V. villosa* chromosome scaffolds led us to question if there was significant divergence in genic sequence of the species. Using a similar strategy to Xi et al. [11], we used the protein coding sequence of nine legume species (Table 6) to estimate gene orthogroups. Orthofinder version 2.5.4 was used to cluster all annotated genes into orthogroups with default parameters [48]. Orthogroup gene assignments were compared across species using the UpSetR version 1.4.0 package (https://cran.r-project.org/web/packages/UpSetR/) in R 4.2.1. Newick files generated by Orthofinder were visualized in the etetoolkit’s “treeview” utility (Figure 8). The Vvill1.0 assembly was found to have the most exclusive orthogroups at 2,555 total orthogroups (Figure 8A). Gene Orthogroup dendrograms (Figure 8B) suggested that gene orthogroup content was similar between the *V.sativa* and Vvill1.0 reference assemblies despite aforementioned structural differences between the two assemblies (Figure 6). We note that this dendrogram does not match the organization of the Fabeae tribe members proposed by Macas et al. (2015). This is mostly due to differences in comparisons between genetic features: where Macas et al. (2015) compared repetitive element conservation and our study compared gene orthogroup sequence conservation. Repetitive elements are often not under selective pressures and are more frequently subject to mutation [49,50] which makes them more informative for comparisons of closely related members of the same species. Comparison of conserved gene orthogroups can better reveal the divergent lineages of different species but such comparisons are only possible after representative genome assemblies have been constructed. Our assembly of the Vvill1.0 reference genome finally allows the accurate placement of *V. villosa* within the Fabeae tribe using conserved gene sequence analysis.

**Table 6.** List of species and their associated genome assemblies used in this study.

| **Species** | **Source of data** | **Version** |
| --- | --- | --- |
| *Vicia villosa* | This project | 1.0 |
| *Vicia sativa* | GigaDB | 1.0 |
| *Vigna unguiculata* | Phytozome | 1.0 |
| *Phaseolus vulgaris* | Phytozome | 2.0 |
| *Lens ervoides* | Phytozome | 1.0 |
| *Lens culinaris* | Phytozome | 2.0 |
| *Medicago truncatula* | INRA | MtA17 r5 |
| *Pisum sativum* | URGI | 1a |
| *Trifolium pratense* | GenBank | 1.1 |

**Figure 8.**
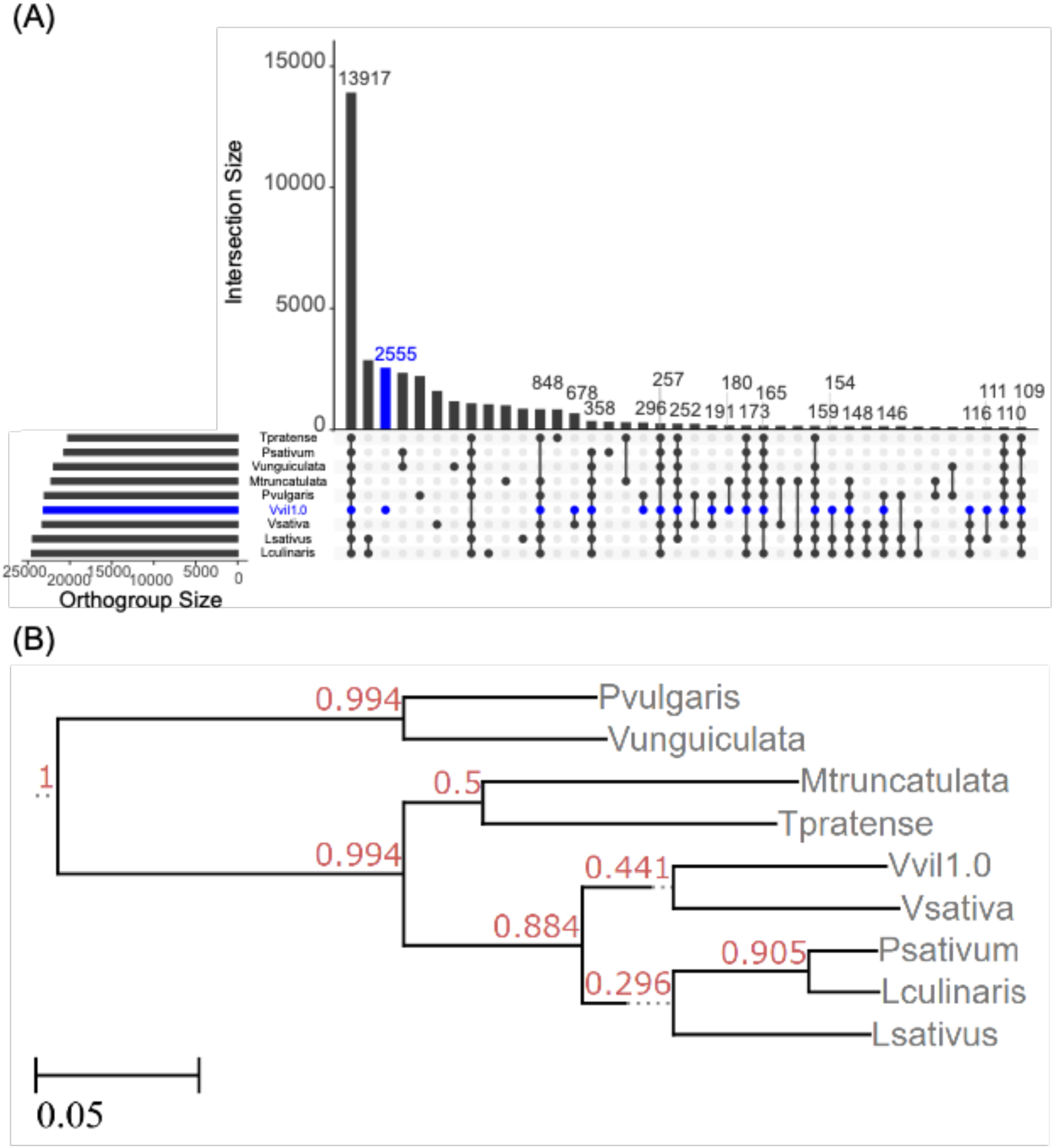
Orthogroup gene comparisons among nine legume species. An upset plot of identified orthogroups (A) suggests that *V. villosa* (blue) has the most unique annotated orthogroups of all compared legume species. Orthogroup dendrogram (B) showing the ortholog-derived relationship of *V. villosa* to other legume species. Values at each node indicate the bootstrap support for each node based on the magnitude of relative error (MRE) test that is the default in the Orthofinder software tool.

## REUSE POTENTIAL

This chromosome-scale genome assembly of *Vicia villosa* provides the foundation for a genetic improvement program for an important cover crop and forage species. Beyond its practical uses, the assembly shows a substantial difference in genome structure compared to a recently released member of the same genus, *Vicia sativa*. These structural differences are in contrast to the conservation of gene orthologs shared by the two species, which suggest that the *V. villosa* assembly may provide an interesting outgroup in comparisons of leguminous plant genomes.

Finally, the documentation of the methods used to resolve a highly heterozygous genome assembly will be useful to resolve issues with assembly of other outcrossing plant species. Specifically, to our knowledge, we are the first to document telomeric “bouquet” patterns during scaffolding using chromatin capture. These methods and our resulting genome assembly will be of use to a wider group of researchers interested in assembling genomes from leguminous plant species.

## AVAILABILITY OF SOURCE CODE AND REQUIREMENTS

The Themis-ASM assembly validation workflow can be found at the following GitHub repository: https://github.com/tdfuller54/Themis-ASM . All other custom scripts used to process the data and generate figures can be found at the following GitHub repository: https://github.com/njdbickhart/ForageAssemblyScripts . A list of software tools and versions used in this analysis is provided in Table 1 of the Supplementary Material.

## DATA AVAILABILITY

All raw sequence data used in the genome assembly and validation can be found in the NCBI’s Sequence Read Archive (SRA) under Bioproject accession PRJNA868110. Genome accession for the Vvill1.0 assembly is under NCBI accession JAROZA000000000. Transcript and other data are available via the Gigascience database, GigaDB.

## AUTHORS’ CONTRIBUTIONS

LMK, TPLS, and MLS generated the genome WGS and Omni-C data. SA generated the transcript sequence data. DMB and TPLS assembled the genome and DMB purged duplicates. HM generated scaffolds from Hi-C read alignments. DMB and TF ran the analysis of the assembly. All authors read and contributed to the final version of the manuscript.

## Supporting information

Uniprot Annotation Data

NCBI-NR Annotation Data

Output from EggNOGmapper

## ACKNOWLEDGEMENTS

We thank Dr. Kristen Kuhn, Kelsey McClure, and Dr. Jennifer McClure for technical assistance and Dr. Maria Monteros for creation of the HV-30 line. This project was supported in part by an appointment (of SA) to the Research Participation Program at the U.S. Dairy Forage Research Center, ARS-USDA, administered by the Oak Ridge Institute for Science and Education through an interagency agreement between the U.S. Department of Energy and ARS-USDA. The USDA does not endorse any products or services. Mentioning of trade names is for information purposes only. The USDA is an equal opportunity employer.

## LIST OF ABBREVIATIONS

bp: base pairs
BUSCO: benchmark universal single-copy ortholog
Gbp: gigabase pairs
kbp: kilobase pairs
KEGG: Kyoto Encyclopedia of Genes and Genomics
KO: KEGG orothologous groups
LINE: long interspersed nuclear element
LTR: long terminal repeat
Mbp: megabase pairs
MITE: miniature inverted-repeat transposable elements
MRE: magnitude of relative error
NCBI: Nation Center for Biotechnology Information
PE: paired end
SRA: Sequence Read Archive
Vvill1.0: *Vicia villosa* reference-quality genome assembly
WGS: whole genome sequencing

## FIGURE LEGENDS

## TABLES

