## Supplementary Material for "A reference assembly for the legume cover crop, hairy vetch (*Vicia villosa*)"

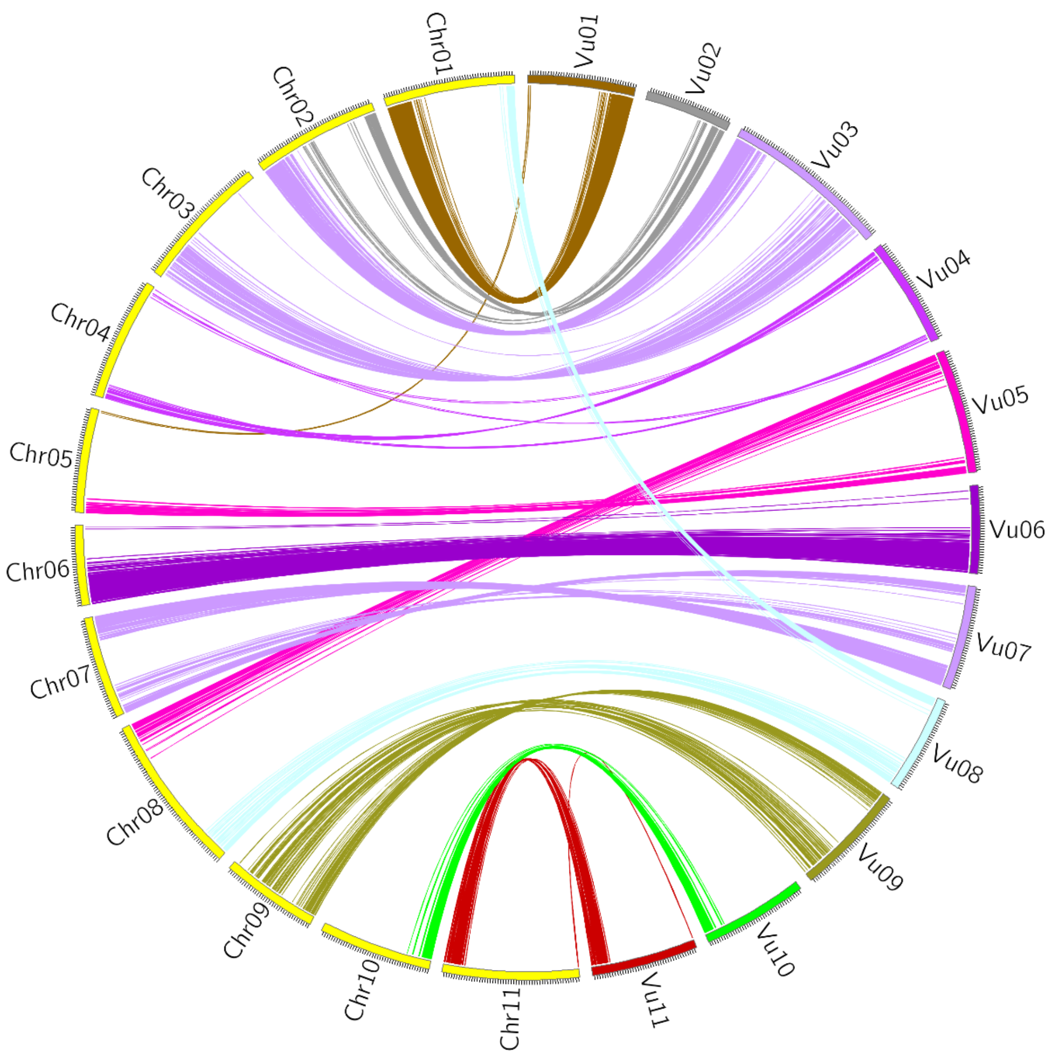


**Supplemental Figure 1.** Circos plot showing syntenic regions shared between *Phaseolus vulgaris* (yellow) and *vigna unguiculata* (multi-colored) genomes [43]. Ribbons (colored matching the *vigna unguiculata* scaffolds) represent pairwise alignments of 100,000 bp or larger identified using minimap2 [42].

Supplemental Table 1. Software and versions used in assembly and analysis of Vvill1.0

| Software | Version |
|---|---|
| BUSCO | 5.3.2 |
| BWA-MEM | 0.7.17-r1188 |
| DIAMOND | 2.0.14.152 |
| EDTA | 2.0.0 |
| EggNOGmapper | 2.1.8 |
| FRC_align | 1.0.0 |
| Freebayes | 1.3.1 |
| GenomeScope | 1.0.0 |
| Jellyfish | 2.2.9 |
| Juicebox | 2.20.00 |
| LUMPY-SV | 0.3.1 |
| merqury | 1.3 |
| meryl | 1.4 |
| minimap2 | 2.24 |
| Orthofinder | 2.5.4 |
| PacBio IPA | 1.3.1 |
| PacBio SMRT Link | 9.0 |
| purge_dups | 1.0.1 |
| RepeatMasker | 4.0.6 |
| RepeatModeler (RRID:SCR_015027) | 2.0.4 |
| SAMBLASTER | 0.1.26 |
| SAMtools | 1.15.1 |
| STAR (RRID:SCR_004463) | 2.7.9 |
| UpSetR | 1.4.0 |
